# Sphingosylphosphorylcholine (SPC) is a substrate for the *Pseudomonas aeruginosa* phospholipase C/sphingomyelinase, PlcH

**DOI:** 10.1101/2025.03.27.645745

**Authors:** Pauline DiGiannivittorio, Kristin Schutz, Lauren A. Hinkel, Matthew J. Wargo

**Affiliations:** Department of Microbiology and Molecular Genetics, Larner College of Medicine, University of Vermont; Cellular, Molecular, and Biomedical Sciences Graduate Program, University of Vermont

**Keywords:** sphingosylphosphocholine, sphingosine, lipid, pathogenesis

## Abstract

Sphingolipids are critical to eukaryotic cell membrane structure and function and play important roles in a variety of host processes that impact infection. Thus, it is not surprising that many pathogens can perturb host sphingolipid homeostasis, often to promote pathogenesis. *Pseudomonas aeruginosa* is a common opportunistic pathogen that, among many virulence factors, secretes the dual-functioning hemolytic phospholipase C/sphingomyelinase, PlcH. PlcH contributes to *P. aeruginosa* pathogenesis in several ways and *plcH* mutants are defective in nearly every infection model, wherein PlcH has been shown to hydrolyze both phosphatidylcholine and sphingomyelin, resulting in inflammation and rupture of host cell membranes. Here, we demonstrate that PlcH can also hydrolyze sphingosylphosphocholine (SPC, also known as lysosphingomyelin), an important host signaling sphingolipid responsible for regulating cellular and tissue responses such as inflammation and endothelial barrier function. PlcH hydrolyzes sphingomyelin to generate phosphocholine and ceramide, and analogously, here we demonstrate that PlcH hydrolyzes SPC to sphingosine and putatively, phosphocholine. We provide evidence that SPC induction of PlcH is primarily regulated by the sphingosine-responsive SphR regulator and that resultant sphingosine liberated from SPC induces transcription from the other genes in the SphR regulon. This work introduces another way that *P. aeruginosa* can alter the host sphingolipidome, potentially a different mechanism to promote pathogenesis. The capacity for the hemolytic *Clostridium perfringens* alpha toxin to also cleave SPC suggests that SPC may be a common substrate for phosphocholine-specific phospholipases C.

**Importance:** PlcH is a secreted phospholipase C/sphingomyelinase that is important for the virulence of *P. aeruginosa*. Here we show that both *P. aeruginosa* PlcH and *C. perfringens* alpha toxin can hydrolyze the signaling phospholipid sphingosylphosphorylcholine (SPC), also called lysosphingomyelin. Thus, SPC should be considered a potential target for such phospholipases during infection whose resulting hydrolysis can induce sphingosine-sensitive genes.

## Introduction

The secreted enzyme PlcH is an important virulence factor for *P. aeruginosa* pathogenesis and infection^1-4^ and has both phosphocholine-specific phospholipase C (PLC) and sphingomyelinase activity^5^. PlcH hydrolyzes phosphatidylcholine into phosphocholine and diacylglycerol and hydrolyzes sphingomyelin into phosphocholine and ceramide^1^, while lipids with other headgroups are very poor PlcH substrates^6,7^. Secreted PlcH, primarily in complex with the chaperone PlcR2^7-10^, causes cellular and tissue damage, promotes inflammation, and disrupts lung surfactant function, enhancing the pathogenesis and *in vivo* survival of *P. aeruginosa*^11-19^. Transcription of *plcH* is controlled by three independent activators, whose inducing signals are present in the host: the glycine betaine and dimethylglycine-responsive transcriptional activator GbdR^20-22^, the sphingosine-binding transcriptional activator SphR^23^, and the phosphate starvation-responsive transcriptional activator PhoB^24^. Choline induces *plcH* transcription and resultant enzyme secretion only after choline metabolism to glycine betaine^21,25^. Phosphatidylcholine and sphingomyelin are considered the primary PlcH substrates in vivo^6,7^, however the ability of PlcH to remove phosphocholine from other molecules, like the colorimetric substrate nitrophenylphosphorylcholine (NPPC)^7,26^, suggest that other phosphocholine-containing molecules in the host might be physiologically relevant PlcH substrates.

Sphingosylphosphocholine (SPC) (**Fig 1**), also called lysosphingomyelin, is a bioactive sphingolipid with functional similarity to sphingosine-1-phosphate (S1P)^27,28^. SPC can be generated through the removal of the N-acyl tail from sphingomyelin via sphingomyelin deacylase and can also be synthesized de novo in specific cell types, such as platelets^29-31^, and is associated with both high-density (HDL) and low-density (LDL) lipoproteins^32^. As a signaling molecule, SPC can bind and activate S1P receptors 1-5^33^ and has also been implicated as a second messenger regulating intracellular Ca^2+^ levels^34-36^. SPC is involved in the maintenance of cell proliferation^37^, differentiation^38^, and regulation of apoptosis^39^, and has impacts on regulation of the immune response^40^ and endothelial barrier function^36,41,42^.

**Figure 1:**
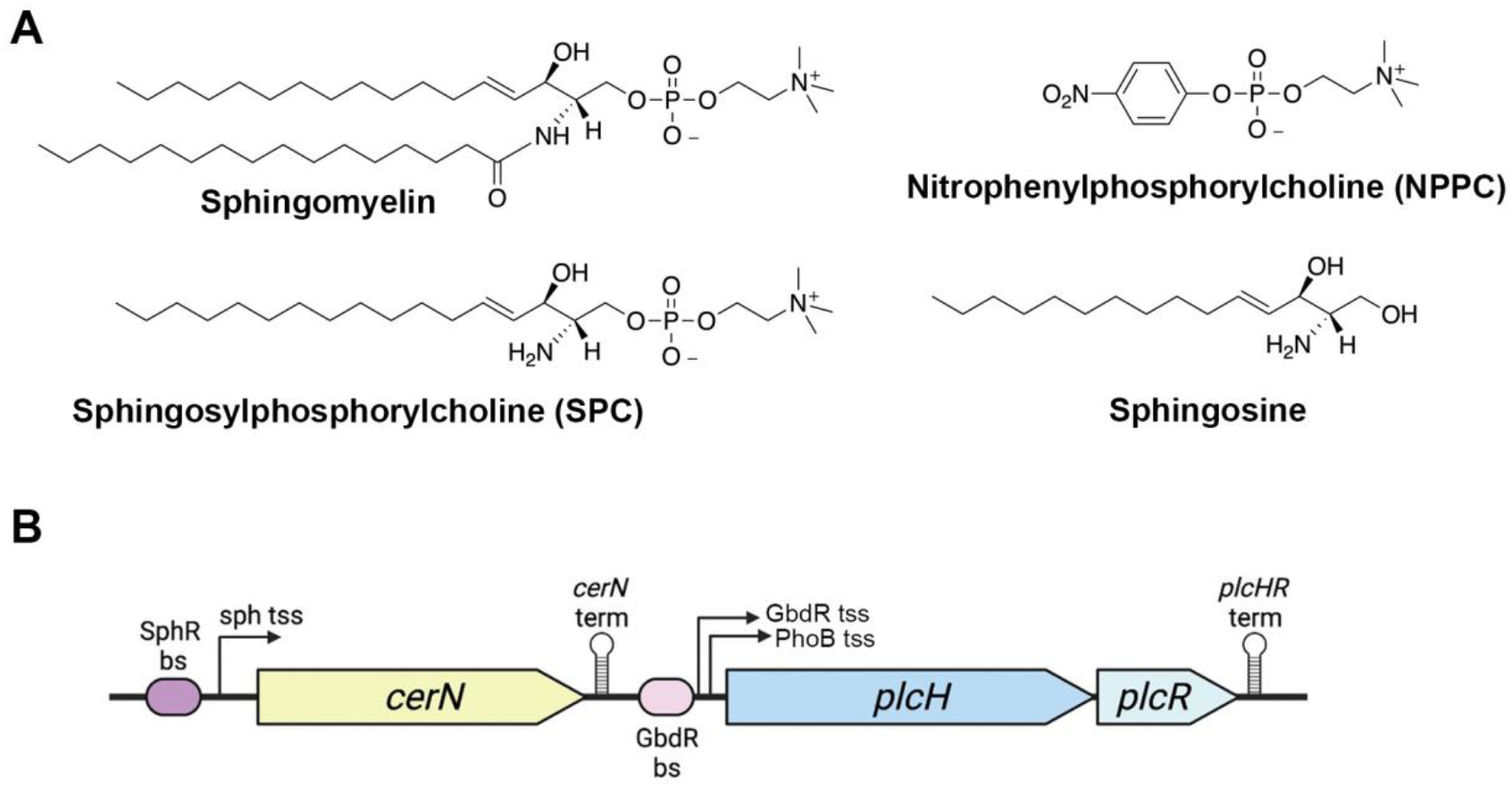
**(A)** Structures of sphingomyelin, SPC, NPPC, and sphingosine and **(B)** Schematic of the *plcH* locus with SphR and GbdR binding sites. Chemical structures generated with ChemDraw and gene organization diagram generated with BioRender. Abbreviations: bs, binding site; sph, sphingosine; tss, transcription start site; term, transcriptional terminator.

Here, we show that SPC is a PlcH substrate and that SPC induces *plcH* transcription and thus, PlcH activity primarily via SphR detection of the sphingosine produced by SPC hydrolysis. Additionally, the purified PLC alpha toxin from *Clostridium perfringens* also hydrolyzes SPC, pointing to SPC as a phosphocholine-PLC substrate more generally. These findings add an important host signaling molecule to the substrate repertoire of PlcH and other bacterial PLCs.

## Results

### SPC induces PlcH expression and transcription of the other SphR-regulon members

Following up on our identification of sphingosine as an inducer of *plcH* transcription via SphR^23^, we first investigated whether SPC could induce PlcH activity, as measured by *p*-nitrophenyl phosphorylcholine (NPPC) hydrolysis. After a 4-hour incubation of PA14 WT with SPC, there was a significant increase in PlcH activity compared to the pyruvate negative control (**Figure 2A**). Sphingosine and choline conditions were included as positive controls, as we have previously shown that sphingosine induces *plcH* in an SphR -dependent manner, whereas choline induces *plcH* in a GbdR-dependent manner^20-22^.

**Figure 2:**
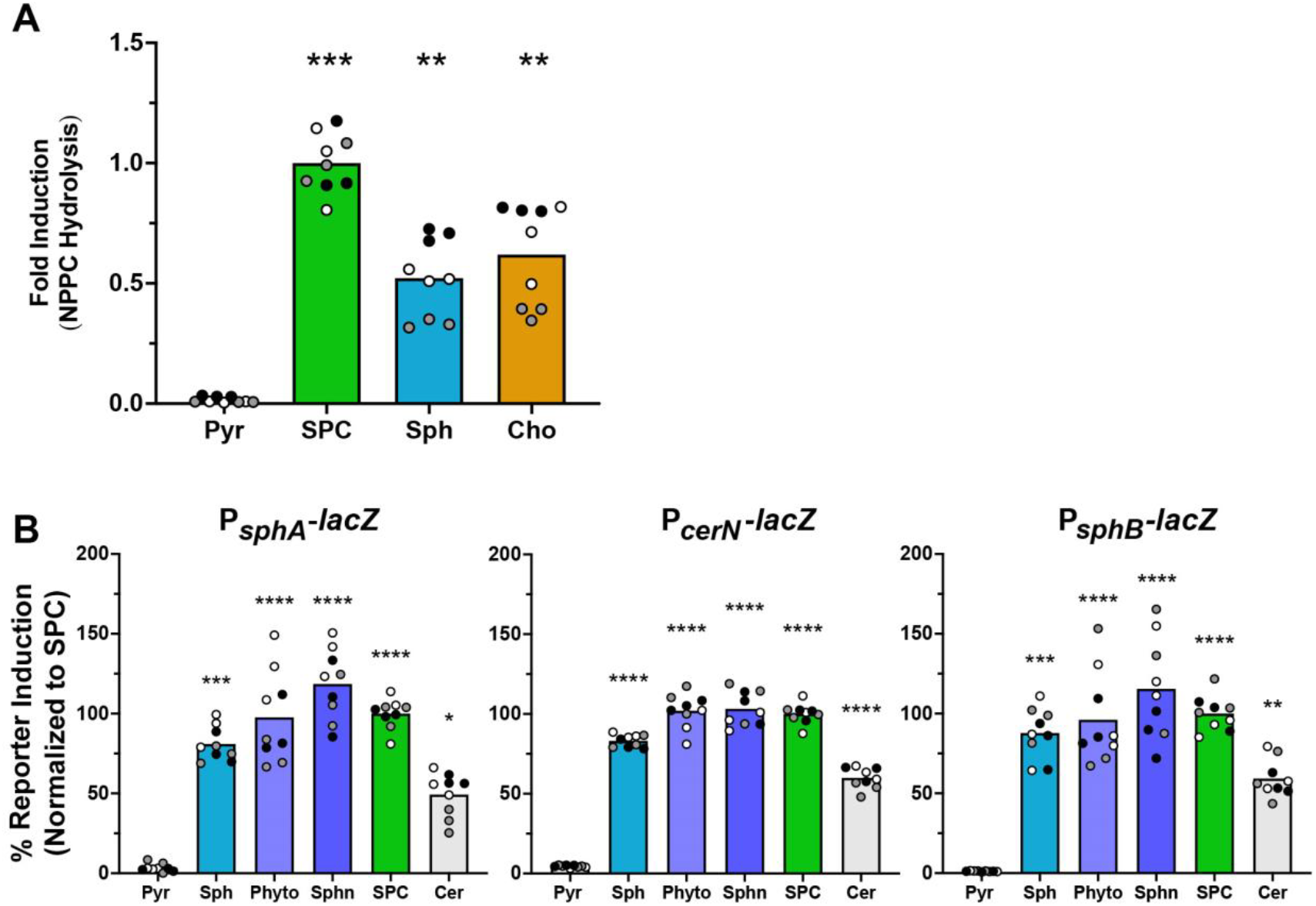
SPC induces PlcH activity and transcription of the SphR regulon. **(A)** SPC induces extracellular PlcH activity as measured by nitrophenylphosphorylcholine (NPPC) hydrolysis as normalized to the level induced by SPC. **(B)** β-galactosidase activity normalized to signal from the SPC induction condition using the plasmid-borne *sphA* promoter *lacZ* reporter (left), the chromosomal *lacZ* integration into the *cerN* locus generating a synthetic *lacZcerN* operon (middle), and the chromosomal *lacZYA* integrated into the *sphBCD* locus replacing the native genes and generating a reporter of the *sphB* promoter (right). Statistical significance for both A & B noted as * (p < 0.05), ** (p < 0.01), *** (p < 0.001), and **** (p < 0.0001) using 1-way ANOVA and Dunnett’s post-test with pyruvate as the comparator. For all panels, all collected data points are shown and are colored by experiment with white circles for all replicates from experiment #1, grey from experiment #2, and black from experiment #3. Only the means for each experiment are used in the statistical analyses for these panels (n = 3 per condition). Abbreviations: Pyr, pyruvate (control); SPC, sphingosylphosphorylcholine; Sph, sphingosine; Sphn, sphinganine; Phyto, phytosphingosine; Cer, ceramide.

Since sphingosine induces expression of *P. aeruginosa* genes involved in ceramide and sphingosine metabolism in addition to *plcH*^23,43^, we next investigated whether SPC induced these same sphingosine-responsive genes. After a 4-hour incubation with SPC, the reporter constructs for the *sphA, cerN*, and *sphB* promoters were each induced, relative to the negative control pyruvate-only condition (**Figure 2B**). Sphingosine and its analogues sphinganine and phytosphingosine were included as positive induction controls^44^, while ceramide was included as a control for a compound that must be metabolized to sphingosine (by the ceramidase, CerN) to allow induction of the sphingosine-responsive genes^43,45^. Given the similarity between SPC induction and sphingosine induction, we predicted that SPC is likely hydrolyzed to sphingosine to enable induction.

### SPC hydrolysis is *plcH* dependent and generates sphingosine

PlcH can hydrolyze sphingomyelin to ceramide and phosphocholine. Considering the structural similarity of SPC to sphingomyelin (**Figure 1A**), we predicted that PlcH would also hydrolyze SPC, generating sphingosine and phosphocholine as products. To test this hypothesis, we first used the *sphA* promoter *lacZ* reporter construct (from **Figure 2B**, left panel) in both *P. aeruginosa* WT and Δ*plcHR*. Upon exposure to SPC, the Δ*plcHR* mutant showed substantially reduced reporter induction compared to WT (**Figure 3A**). Sphingosine induction was not different between these strains. These data supported a role for PlcH in SPC hydrolysis.

**Figure 3:**
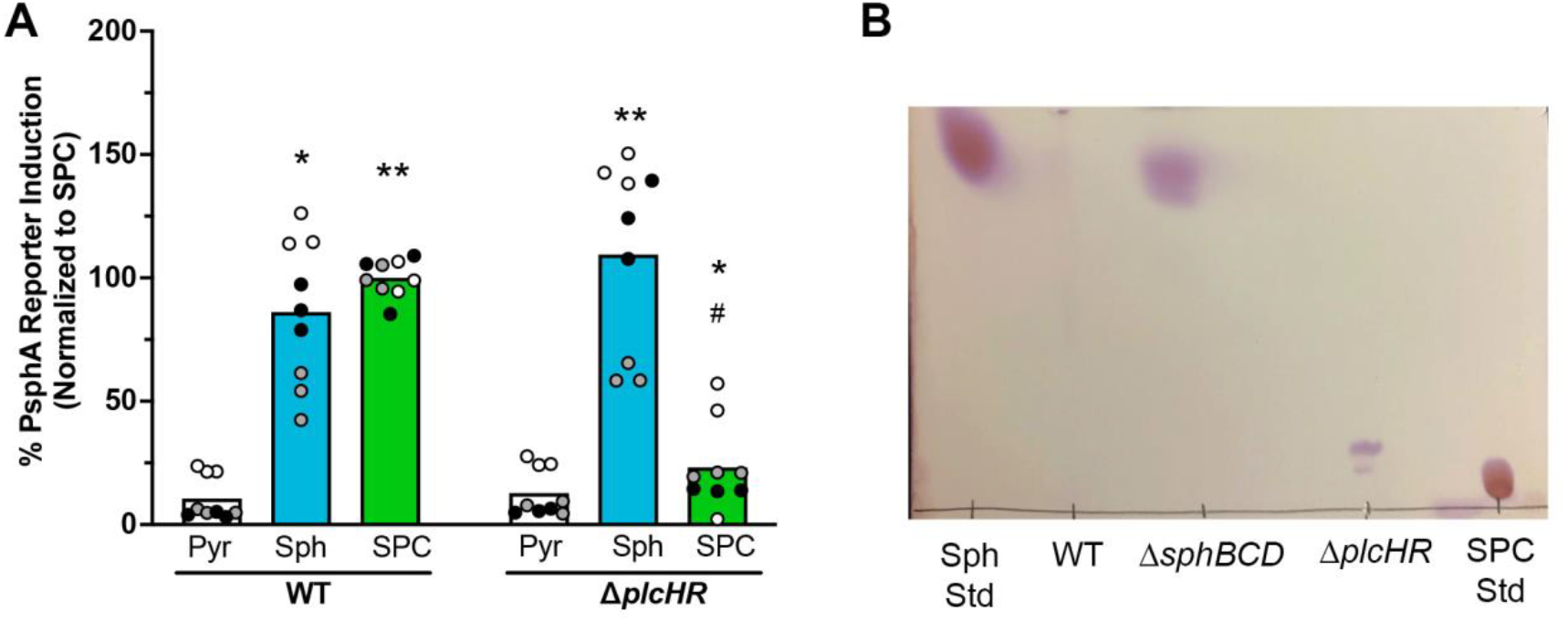
*plcH* is necessary for SPC hydrolysis to sphingosine. **(A)** WT carrying the P_*sphA*_*-lacZ* reporter responds to SPC, while Δ*plcHR* carrying the same reporter does not. **(B)** Using thin layer chromatography of lipid extracts from strains supplied with SPC, when SPC is supplied to the Δ*plcHR* strain, sphingosine is not generated (see migration of the standards on either side of the TLC) . TLC was done on silica plates with a chloroform:methanol:water (65:25:4) mobile phase and after drying, were stained with ninhydrin to detect the primary amine on the sphingoid base. Data shown as means of three independent experiments. Statistical significance noted as * or ^#^ (p < 0.05) and ** (p < 0.01) using 2-way ANOVA with Sidak’s post-test showing comparisons within reporter strain compared to the pyruvate condition (asterisks) or comparing WT + SPC to Δ*plcHR* + SPC (#). For (A), all collected data points are shown and are colored by experiment with white circles for all replicates from experiment #1, grey from experiment #2, and black from experiment #3. Only the means for each experiment are used in the statistical analyses for these panels (n = 3 per condition). Abbreviations: SPC, sphingosylphosphorylcholine; Sph, sphingosine; Pyr, pyruvate; Std, standard.

The sphingosine-sensitive reporter assay is an indirect method to assess sphingosine production, so to directly visualize the sphingosine formed upon SPC hydrolysis by PlcH we performed thin layer chromatography (TLC). Lipids extracted from *P. aeruginosa* Δ*sphBCD* supernatants exposed to SPC show sphingosine formation, while no sphingosine is seen in extracts from Δ*plcHR* exposed to SPC (**Figure 3B**). No sphingosine remains in the WT supernatants due to sphingosine metabolism which requires *sphB* and *sphC*, as we have recently shown^44^.

### The *Clostridium perfringens* alpha toxin, a hemolytic phospholipase C, also hydrolyzes SPC to generate sphingosine

The alpha toxin from *C. perfringens* is a hemolytic phospholipase C with specificity for phosphocholine-containing lipids (abbreviated *Cp* PLC). As we have yet to purify active PlcH, we tested whether purified *Cp* PLC was capable of SPC hydrolysis. Pretreatment of SPC with purified *Cp* PLC led to reporter induction from the *plcHR* deletion strain carrying the chromosomal sphingosine-responsive reporter (*lacZ* at the *cerN* locus), while incubation of SPC with the buffer control led to no induction in the *plcHR* deletion strain (**Figure 4A**). These data supported the ability of *Cp* PLC to hydrolyze SPC. Using thin-layer chromatography, we also demonstrate complete conversion of SPC to sphingosine within our limit of detection (**Figure 4B**). Thus, SPC is likely a general substrate for PLCs that prefer phosphorylcholine headgroups.

**Figure 4:**
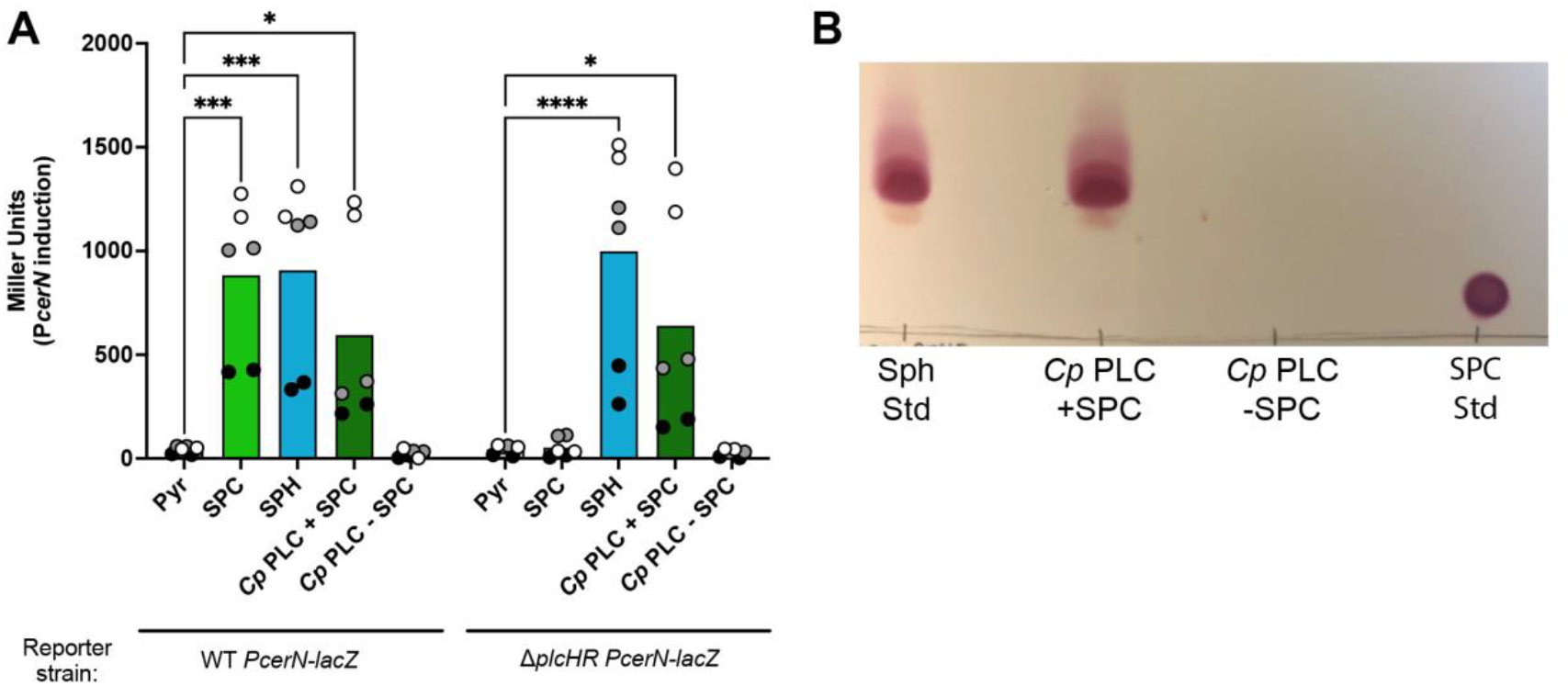
The *Clostridium perfringens* alpha toxin hydrolyzes SPC to generate sphingosine. **(A)** Wild type with a chromosomal P_*cerN*_*-lacZ* reporter (same reporter as in Fig 2B, center panel) responds to SPC, while the same reporter in Δ*plcHR* does not. However, treatment of SPC with purified *C. perfringens* PLC enables responsiveness from the Δ*plcHR* reporter strain. **(B)** Using thin layer chromatography, when SPC is incubated with *Cp* PLC sphingosine is generated (see migration of the standards on either side of the TLC). TLC is stained with ninhydrin to detect the primary amine on the sphingoid base. Statistical significance for (A) noted as * (p < 0.05), *** (p < 0.001), and **** (p < 0.0001) using using 2-way ANOVA with Sidak’s post-test with pyruvate as the comparator within each strain. For (A), all collected data points are shown and are colored by experiment with white circles for all replicates from experiment #1, grey from experiment #2, and black from experiment #3. Only the means for each experiment are used in the statistical analyses for these panels (n = 3 per condition). Abbreviations: Pyr, pyruvate (control); SPC, sphingosylphosphorylcholine; SPH or Sph, sphingosine; *Cp* PLC, *Clostridium perfringens* phospholipase C; Std, standard.

### The roles for SphR and GbdR in PlcH induction by SPC

PlcH hydrolysis of SPC hydrolysis results in production of a sphingoid base (**Figure 3B**) and, very likely, phosphocholine, both of which could independently lead to induction of *plcH* transcription. We thus tested whether PlcH induction by SPC was regulated by SphR, GbdR, or a combination. Upon exposure to SPC, we measured PlcH activity by NPPC hydrolysis in WT, Δ*sphR*, Δ*gbdR*, and engineered strains with mutations in the SphR binding site of the *cerN* promoter, mutations in the GbdR binding site of the *plcH* promoter, and a strain containing both SphR and GbdR binding site mutations. While SPC induced PlcH enzyme activity in WT, the Δ*sphR* mutant and the strain with mutation of the SphR binding site showed no induction in the presence of SPC (**Figure 5**). Interestingly, the Δ*gbdR* and GbdR binding site mutants showed increased PlcH activity compared to WT when exposed to sphingosine but not SPC (**Figure 5**). These data indicate that PlcH induction by SPC is under SphR transcriptional control at this tested concentration of SPC. The SphR and GbdR binding site double mutant strain functioned as our negative control, since neither product of SPC hydrolysis would be capable of inducing PlcH, which also provides support that there is no way for SPC to induce PlcH in the absence of its hydrolysis to phosphocholine and sphingosine. Sphingosine and choline conditions were included as positive controls for each transcriptional regulator system.

**Figure 5:**
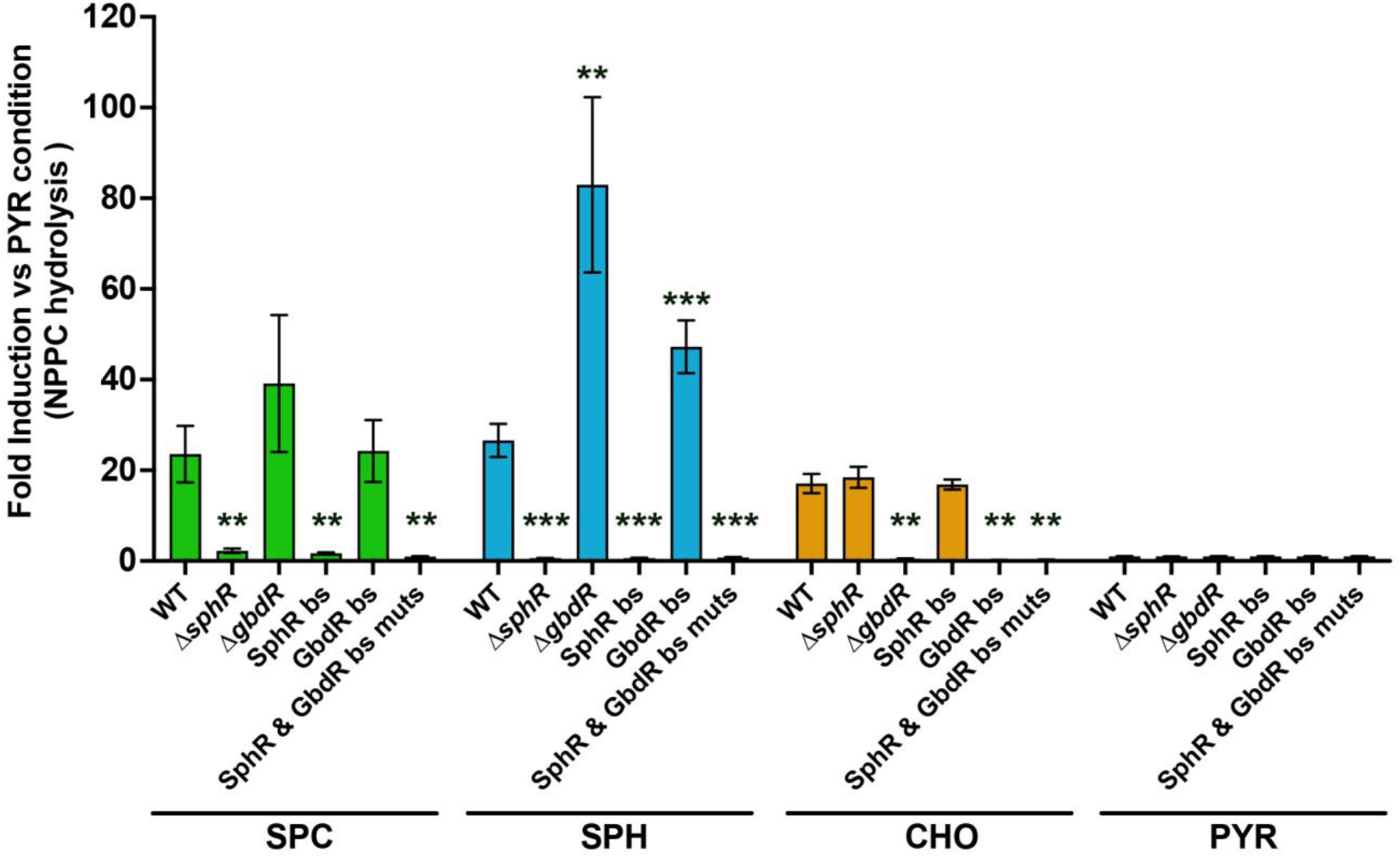
Contributions of GbdR- and SphR-dependent regulation to SPC induction of secreted PlcH enzyme activity. Secreted PlcH enzyme activity measured by NPPC hydrolysis and normalized by fold induction to the pyruvate condition for each strain. Data shown as means of three independent experiments with standard error bars, as the number of groups made plotting all data points, as done for the other figures, impractical. Statistical significance noted as * (p < 0.05), ** (p < 0.01), and *** (p < 0.001) using 2-way ANOVA with Sidak’s post-test showing comparisons within induction condition to the WT strain. Abbreviations: bs, binding site; mut, mutant; SPC, sphingosylphosphorylcholine; SPH, sphingosine; CHO, choline; PYR, pyruvate.

## Discussion

Here, we show that the *P. aeruginosa* virulence factor PlcH can hydrolyze SPC resulting in sphingosine production. This finding suggests that, in addition to the classical substrates phosphatidylcholine and sphingomyelin considered during infection, *P. aeruginosa* may also be capable of perturbing host signaling via SPC hydrolysis. Since the *C. perfringens* PLC also shows SPC hydrolysis, this suggests that SPC may be a target for other phosphocholine-specific PLCs. SPC is structurally similar to sphingomyelin (**Fig. 1a**) and PlcH can hydrolyze a range of phosphocholine-containing compounds, therefore it is not surprising that SPC hydrolysis by *P. aeruginosa* is dependent on PlcH. The hydrolysis of SPC by both *P. aeruginosa* and *C. perfringens* PLCs, which are very different by sequence and structure^46-48^, supports the idea that phosphocholine recognition is the primary driver of hydrolysis by these enzymes. While this has been well studied for the phosphocholine-hydrolyzing PLCs of *C. perfringens* and *B. cereus*, substrate recognition by PlcH is not well-described. Apart from the active site threonine (T178), discovered by homology to *Francisella tularensis* AcpA and subsequently experimentally tested^7,9,19^, very little is known about PlcH recognition of the phosphocholine headgroup or the moiety attached to the phosphocholine. Since the affinity of PlcH for NPPC is much lower than for phosphatidylcholine or sphingomyelin^9,19^, some portion of the acyl moieties on these molecules are likely recognized.

SPC can induce PlcH production, and it does so primarily through SphR-dependent induction, a pathway we have recently described^23^. The primacy of the sphingosine moiety for PlcH induction may have more to do with the SPC concentration used during these experiments than it does the comparative importance of the individual regulators per se. The responsiveness of GbdR to GB produced from exogenous choline is less sensitive than SphR detection of exogenous sphingosine. SphR can detect exogenous sphingosine as low as 2.5 µM with a maximal response at 200 µM^23,43,44^, whereas the lower limit for choline detection is ∼3 µM ^49,50^, governed by the transport Kd, with a maximal response at 2 mM ^24^. Thus, at the concentration used in these experiments and the experimental timing, the sphingosine released from SPC hydrolysis is more important for PlcH induction than the phosphocholine. This relationship might not be the same at all concentrations, time steps, or in vivo.

The data presented here use induction conditions with different concentrations of SPC. To measure PlcH activity, *P. aeruginosa* strains were induced with 100 μM SPC, while for reporter assays, we used 20 μM SPC. Within the human body, SPC is commonly seen at a concentration estimated around 50 ± 15 nM, a concentration much lower than the concentrations tested in this study^30,51^. However, steady state levels in a whole compartment (like the blood) are often much lower than concentrations within local environments in which a product is being actively produced, such as in association with platelets. Thus, it remains an open question whether PlcH hydrolysis of SPC happens in vivo and whether *P. aeruginosa* SPC hydrolysis has any impact on the host during infection. Given SPC’s important roles in regulating endothelial cell and barrier function, future studies should investigate alterations in these responses and their alteration in response to WT and *plcH* mutant strains.

## Materials and Methods

### Strains and growth conditions

*Pseudomonas aeruginosa* PA14 and isogenic mutant strains (**Table 1**) were maintained on Lysogeny Broth-Lennox formulation (LB) or *Pseudomonas* isolation agar (PIA) plates with 50 µg/mL gentamicin added when appropriate. *Escherichia coli* strains used in this study were maintained on LB plates or liquid LB supplemented with 10 µg/mL or 7 µg/mL gentamicin, respectively. During genetic manipulations, *P. aeruginosa* was selected for, and *E. coli* selected against, using PIA plates supplemented with 50 µg/mL gentamicin. Prior to transcriptional and enzyme induction studies, *P. aeruginosa* was grown at 37°C overnight in morpholinepropanesulfonic acid (MOPS) medium^52^ modified as previously described^53^, and supplemented with 20 mM pyruvate and 5 mM glucose.

**Table 1:**
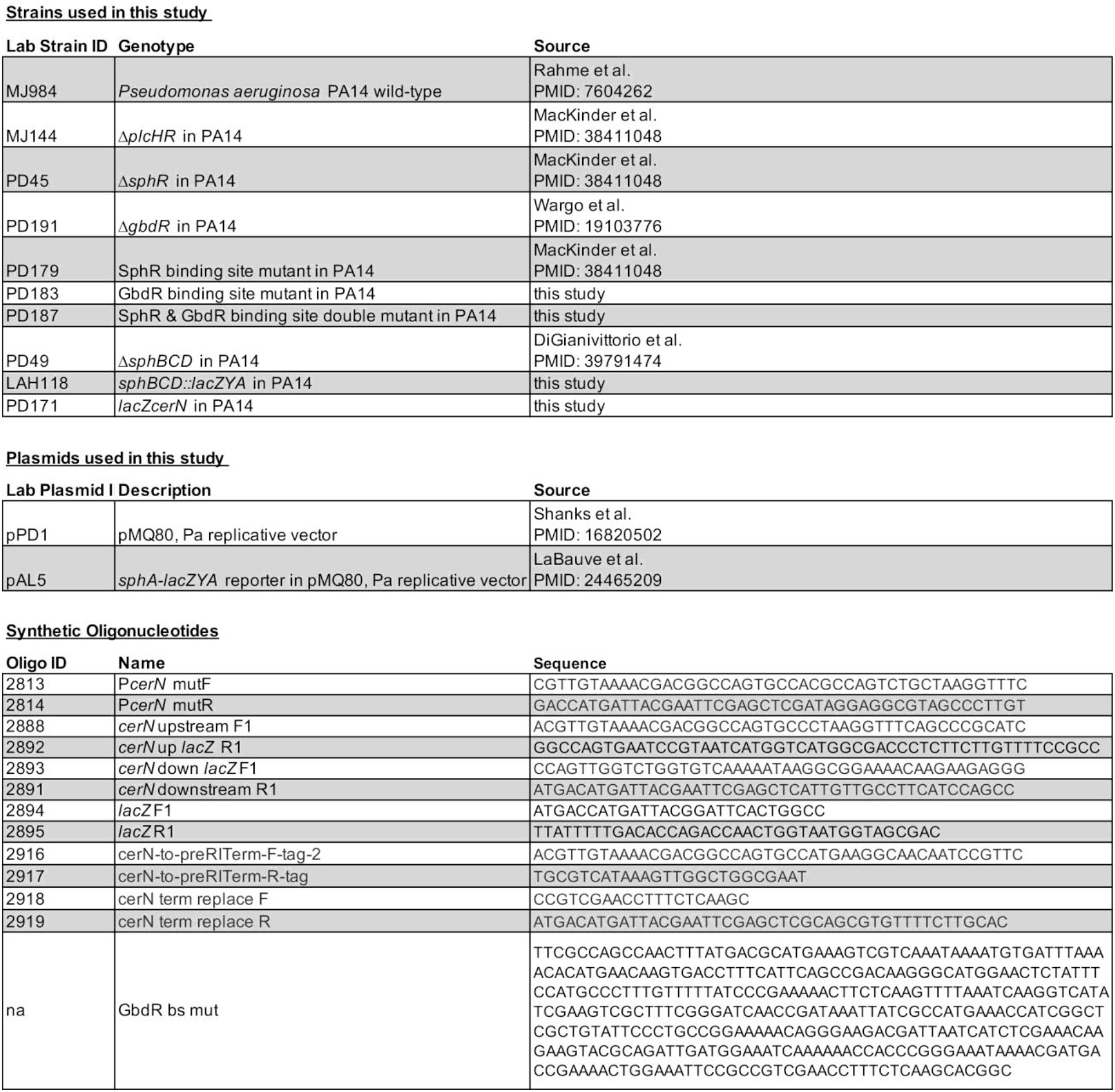
Strains, plasmids, and oligonucleotides used in this study.

### General Allelic Exchange, Chromosomal Alterations, and Electroshock Transformations

All allelic exchange constructs were generated using the pMQ30 non-replicative, counter-selectable vector^54^. Briefly, after constructs were cloned into the pMQ30 backbone, they were transformed into chemically competent S17 λ*pir E. coli*. For conjugation, donor *E. coli* were mixed with respective recipient *P. aeruginosa* strains, pelleted via centrifugation, resuspended in a small volume of LB, and spotted onto LB plates, and incubated overnight at 30 °C. Single-crossover integrants were selected by plating on PIA with 50 μg/ml gentamicin following incubation at 37 °C for 24 hours. Selected single crossover integrants were inoculated into LB, incubated at 37 °C for 3-4 hours with shaking, and plated onto LB and LB with no NaCl containing 5 % sucrose and incubated overnight at 30 °C. Sucrose resistant colonies were screened for loss of gentamicin resistance prior to PCR screening to determine whether each double-crossover colony was a mutant or WT revertant.

Briefly, the allelic exchange vector for mutation of the GbdR binding site in the *plcH* promoter used HiFi assembly (NEB) of two PCR products generated from PA14 genomic DNA (using primers 2916 & 2917 for the upstream side and 2918 & 2919 for the downstream side), a synthetic fragment containing the GbdR binding site mutation region “GbdR bs mut”, and HindIII+KpnI cut pMQ30. Sequence verified plasmids were transformed into chemically competent S17λ*pir E. coli* and allelic exchange using recipient PA14 strains was completed as described above, resulting in strains PD183 (GbdR bs mutant) and PD187 (SphR bs & GbdR bs double mutant). Generation of the SphR binding site mutant in the *cerN* promoter was previously described^23^.

The allelic exchange vector for generation of the *lacZcerN* synthetic operon at the chromosomal *cerN* locus in PA14 was built using HiFi assembly (NEB) from three PCR products (amplifying *cerN* upstream and downstream fragments and *lacZ* fragment) and HindIII+KpnI cut pMQ30. The *cerN* upstream fragment was amplified with primers #2888 & #2892 and the downstream region using #2893 & #2891, both using PA14 genomic DNA as the template. The *lacZ* gene was amplified from pMW5^20^ with primers #2894 and #2895. Sequence verified plasmids were transformed into chemically competent S17λ*pir E*.*coli* and allelic exchange was completed as described above, resulting in strain PD171 (*lacZ-cerN*).

The allelic exchange vector for P_*sphB*_*-lacZYA* incorporation into the PA14 chromosome was built by amplifying the region upstream of the *sphBCD* operon from *P. aeruginosa* PAO1 using primers #2082 and #2286, digestion of the product with HindIII and KpnI, and ligation into similarly cut pGW78, resulting in interim plasmid 1. The downstream region of the *sphBCD* operon was amplified with primers #2287 and #2288, digestion with enzymes NheI and SphI and ligation into similarly cut interim plasmid 1 at the 3’ end of *lacZYA*, yielding interim plasmid 2. The *lacZYA* with *sphBCD* flanking regions was cut from interim plasmid 2 with HindIII and SphI and ligated into similarly cut pMQ30 yielding plasmid p*lacZYA::sphBCD*. Conjugation and allelic exchange were conducted as described above, resulting in strain LAH118.

### Chemicals and notes on sphingolipid stability, solubility, and handling

All media, media components, and standard chemicals were purchased from Thermofisher or Sigma. Sphingolipids such as sphingosine, phytosphingosine, sphinganine, sphingosylphosphocholine, and ceramide were purchased from either Cayman Chemicals or Avanti Polar Lipids. All sphingolipids were dissolved in 95% ethanol (with sonication when necessary) and stored as 50 mM stocks at -20°C. The storage of sphingolipids in aliquot form is critical, as multiple freeze-thaw cycles lead to decrease in sphingolipid potency and function (i.e., antimicrobial activity for sphingoid bases and the ability to stimulate gene induction via SphR^43,44^). Sphingolipids were delivered to culture vessel in ethanol and ethanol was evaporated either by air drying or a gentle stream of nitrogen gas.

### Phospholipase C activity assays (NPPC assays)

As a readout for PlcH activity (i.e., phospholipase C activity), we measured hydrolysis of the synthetic substrate *p*-nitrophenylphosphorylcholine (NPPC) based on the methodology of Kurioka and Mastuda^44^ and modified as previously described^20^. Briefly, *P. aeruginosa* strains were grown overnight shaking at 37°C in MOPS media with 25 mM pyruvate and 5 mM glucose, collected by centrifugation, and washed in MOPS media prior to resuspension in MOPS with 25 mM pyruvate. Culture density was adjusted to OD_600_ = 0.5 with MOPS media with 25 mM pyruvate and with or without 100 μM SPC, sphingosine, or 2 mM choline. Cultures were incubated for 4 hours shaking at 37°C. To measure NPPC hydrolysis, one volume of culture was mixed with one volume of 2 X NPPC reaction buffer (200 mM Tris pH 7.2, 50% glycerol, 20 mM NPPC). NPPC hydrolysis was then measured by quantifying absorbance at 410 nm every five minutes for thirty minutes. Prior to normalization, phospholipase C activity was first calculated to determine micromoles of *p*-nitrophenol generated per minute of reaction per optical density (OD_600_), using the nitrophenol extinction coefficient of 17,700 M-1 cm-1^55^.

### *Clostridium perfringens* alpha toxin reaction

To assess if *Clostridium perfringens* alpha toxin hydrolyzes SPC to sphingosine, a bioassay and thin layer chromatography, 2 mg/mL *C. perfringens* alpha toxin (in molecular biology grade water) was incubated with 100 µM SPC for 4 hours at 37°C, shaking. After incubations, lipids were extracted using the Bligh and Dyer method^56^ and were prepared as described in the TLC section.

### Thin layer chromatography (TLC)

To visually assess SPC hydrolysis to sphingosine, we used thin later chromatography. *P. aeruginosa* strains were grown overnight at 37 °C, shaking in MOPS media with 25 mM pyruvate and 5 mM glucose. Cells were collected via centrifugation, washed in MOPS media, and cell pellets resuspended in MOPS media with 25 mM pyruvate. Cultures were adjusted to OD_600_ = 0.5 in MOPS media with 25 mM pyruvate in a mutli-well plate and choline was added to a concentration of 2 mM. Cultures were incubated shaking for 4 hours at 37°C. After inductions, supernatants were moved to 13 x 100 mm borosilicate glass tubes and incubated with 100 μM SPC for 4 hours, shaking at 37°C. After induction period, lipids were extracted using the Bligh and Dyer method^56^. Briefly, chloroform:methanol (1:2; v:v) was added, samples were vortexed, and one volume of water was added. After briefly vortexing, samples were centrifuged for 10 minutes at 14,000 x g. After centrifugation, the lower organic fraction was collected and dried using N_2_ gas before final resuspension in 20 ml of ethanol. TLC silica gel 60 F254 plates (Sigma Aldrich) were pre-run with acetone and allowed to dry. Lipid extracts or standards at 100 µM were spotted onto the silica plates. After all samples were dried, plates were run with a chloroform:methanol:water (65:25:4) mobile phase. After the mobile phase approached the top of the plate, the plate was removed, dried, and sprayed with 0.2% ninhydrin solution (Acros Organics) to detect the sphingolipids by the primary amine group.

### *sphAlacZ, sphB-lacZ*, and *lacZ-cerN* reporter assays

To investigate sphingosine product formation upon SPC hydrolysis, *sphA, sphB*, and *cerN* transcriptional induction was measure using P*sphA-lacZ* (previously described^43,44^), *sphBCD::lacZYA*, and *lacZcerN* reporter strains and constructs, the latter two constructed as described above. *P. aeruginosa* strains were grown overnight at 37°C, shaking, in MOPS media with 25 mM pyruvate, 5 mM glucose, and 20 μg/mL gentamicin when necessary. Cells were collected via centrifugation, washed in MOPS media, and cell pellets were resuspended in MOPS media with 25 mM pyruvate (supplemented with 20 μg/mL gentamicin if necessary). Culture densities were adjusted to OD_600_ = 0.5 in MOPS media with 25 mM pyruvate in multi-well plates with or without 20 μM SPC, sphingosine, sphinganine, phytosphingosine, or ceramide. Cultures were incubated with shaking at 37°C for 4 hours. β-galactosidase assays were performed as previously described^43,44^ using Miller’s method^57^.

## Acknowledgements

We’d like to thank Jacob Mackinder, Jon Boyson, John Barlow, Bruno Martorelli di Genova, and Markus Thali for helpful discussions. We’d also like to thank Rob Hondal for assistance with ChemDraw.

## Funding

NIH NIAID R56AI173006 (MJW)

CYSTIC FIBROSIS FOUNDATION WARGO24G0 (MJW)

NIH NHLBI T32 HL076122 (PD)

NIH NIAID T32 AI055402 (LAH)

